# Hyperbolic Wavelet Interaction Network (HyWinNet): Multi-Scale Geometric Learning for Interpretable Protein-Protein Interaction Prediction

**DOI:** 10.1101/2025.09.01.673587

**Authors:** Qingzhi Yu, Shuai Yan, Wenfeng Dai, Zhengrong Xi, Yuxin Cheng, Xiang Cheng

## Abstract

Understanding the multi-scale organization of protein-protein interactions (PPIs) is fundamental to deciphering cellular signaling, allosteric regulation, and disease mechanisms, yet existing computational approaches fail to simultaneously resolve atomic-scale binding interfaces and pathway-level coordination. We present HyWinNet, a geometric deep learning framework that unifies Lorentzian hyperbolic graph neural networks with spectral graph wavelet transforms to intrinsically model the hierarchical architecture of biological interactions. The core innovation lies in three synergistic components: (1) Hyperbolic embeddings that preserve evolutionary-conserved topological relationships through negatively curved space projections, capturing scale-free properties of protein networks; (2) Multi-scale wavelet decomposition operating at biologically grounded resolutions to jointly analyze local residue contacts and global pathway dynamics; (3) Contrastive learning with stochastic feature dropout that mimics natural binding interface plasticity. Benchmarked against state-of-the-art methods, HyWinNet demonstrates superior performance in recovering known interactions while predicting previously unannotated functional sites validated through independent structural and biochemical studies. By bridging hyperbolic geometry with graph signal processing, this work establishes a new paradigm for analyzing multi-scale biological networks, offering both interpretable computational predictions and testable hypotheses for experimental validation. The framework’s ability to map hierarchical relationships from atomic details to system-level modules provides a transformative tool for drug discovery and mechanistic studies of complex diseases.

## 1. Introduction

Protein-protein interactions (PPIs), one of the core mechanisms of cellular life activities[1], mediate the specific binding and functional coordination between proteins, playing a pivotal role in key biological processes such as signal transduction[2], metabolic regulation, gene expression, immune response, and cell cycle control[3]. At the molecular level, PPIs serve as the structural basis for enzymatic reactions, receptor-ligand recognition, and complex assembly, while also acting as critical nodes in dynamic cellular regulatory networks[4]. For instance, the cooperative binding of transcription factors regulates gene expression[5], and the interaction between kinases and their substrates determines the activation threshold of signaling pathways[6]. In pathological contexts, aberrant PPIs are closely associated with the development of diseases such as cancer, neurodegenerative disorders, and infectious diseases[7]. For example,dysregulation of the interaction between p53 and MDM2 can lead to the loss of tumor-suppressive function[8]. In contrast, the interaction between viral proteins and host proteins is a key strategy for pathogen invasion[9]. Accurately identifying and understanding these interaction sites is essential for elucidating protein functions, disease mechanisms, drug development, and constructing biological network models[10]. Traditional experimental methods like yeast two-hybrid and Co-IP[11, 12] rely on known protein information (e.g., antibodies, libraries), making it difficult to discover novel or uncharacterized interactions, and are often resource-intensive and unable to provide systematic predictions[13]. In contrast, deep learning-based methods offer lower experimental costs and can integrate multi-source information such as sequence, structure, and function for systematic prediction[14, 15].

Deep learning-based methods for predicting Protein-Protein Interaction Sites (PPIS) can generally be categorized into three types[16]. The first is sequence-based methods, which directly predict interactions using protein amino acid sequences without requiring structural information. However, due to the limited information provided by protein sequences, the prediction results are not always ideal[17]. The second type is structure-based methods, which leverage three-dimensional structural information to model binding interfaces or spatial complementarity. These methods explore potential PPIs by exploiting known protein interaction networks[18]. The third category is multi-modal fusion methods, which integrate multi-source data such as sequences, structures, expression profiles, and domain information. By combining sequence and structural insights, this approach aligns more closely with biological principles[19].

Most deep learning-based methods for Protein-Protein Interaction Site (PPIS) prediction employ core architectures such as Convolutional Neural Networks (CNNs)[20], primarily applied to sequence data for extracting local features through convolutional layers, and Graph Convolutional Networks (GCNs), predominantly used for structural data to capture topological information in protein structures via graph convolutional layers[21]. However, existing computational approaches exhibit significant limitations in modeling hierarchical geometric relationships between proteins and their multiscale dynamic interaction patterns[22, 23].

Specifically, traditional GCNs rely on graph neural networks in Euclidean space, where the volume of a sphere grows polynomially with its radius. This property inherently fails to represent scale-free characteristics (exponentially growing connectivity) and hierarchical architectures prevalent in biological networks. Consequently, graph node vectors embedded in Euclidean space suffer from substantial structural distortion, particularly for complex networks with hierarchical or tree-like topologies[24]. Additionally, conventional Graph Neural Networks (GNNs) face challenges in handling large-scale graphs due to the “over-squashing” phenomenon, where inefficient message passing hinders the modeling of long-range node dependencies[25].

To address these challenges, we propose **HyWinNet**, which introduces three key innovations. (1) We design a **Lorentzian hyperbolic graph neural network encoder** that effectively captures the hierarchical structure of biomolecular networks. By embedding graphs into hyperbolic space, HyWinNet models the exponential expansion and scale-free nature of biological systems, preserving topological properties such as power-law degree distributions and small-world characteristics. Additionally, it leverages geometric flows in hyperbolic space to enable continuous-time graph learning with improved computational efficiency. (2) We develop a **multi-view contrastive learning framework** with 20% feature dropout to enhance representation learning. This framework promotes discriminative embeddings by maximizing agreement between different augmented views of the same protein while minimizing similarity with unrelated proteins. (3) We introduce a **multi-scale Graph Wavelet Transform (GWT)** based on symmetric normalized matrix to simultaneously capture local binding patterns and global pathway interactions across four biologically meaningful scales. This hierarchical analysis alleviates the over-squashing problem and enables HyWinNet to reveal both fine-grained molecular interactions and coarse-scale modular structures, offering a more comprehensive and biologically relevant representation compared to single-scale methods.

## 2. Materials and Methods

In this section, we present the utilized dataset and derive the protein-protein relationship matrix from the dataset. Subsequently, based on this relationship matrix, we propose a deep learning model named HyWinNet for predicting protein-protein interactions. The model employs the following innovative framework: First, it projects protein node features into hyperbolic space to capture hierarchical relationships better. Next, it performs Lorentz graph convolution operations in hyperbolic space to enhance feature propagation. Then, multi-view contrastive learning is applied to maximize the discrimination between positive and negative samples. Finally, multi-scale diffusion wavelet transformation is utilized to extract local and global structural information. Eventually, it predicts protein-protein interactions.

### 2.1. Dataset

We utilize two publicly accessible heterogeneous network datasets introduced by Luo et al.[26] and Zeng et al.[27] for our experiments. Following the removal of duplicated and isolated nodes, the experimental data for HyWinNet are prepared. For experimental validation, we adopted an 85-5-10 split ratio, allocating 85% of samples for training, 5% for validation, and 10% for testing. This partitioning strategy ensures sufficient training data while retaining statistically representative validation and test sets.

### 2.2. HyWinNet Model

a. Overall Architecture As depicted in Fig.1, the HyWinNet framework comprises three sequentially integrated processing stages designed to learn hierarchical and multi-scale representations for PPI prediction:
  - Lorentz Hyperbolic Graph Neural Network Encoder (Section 2.2.1): This module processes the input protein feature matrix and adjacency matrix. It first projects the Euclidean features into Lorentzian hyperbolic space via exponential mapping. Subsequently, it performs Lorentz graph convolutions (Eq. 3-6) utilizing hyperbolic activation functions (Eq. 5) to generate hierarchical node embeddings that intrinsically capture biomolecular relationships.
  - Multi-View Contrastive Learning Module (Section 2.2.3): Operating on the hyperbolic node embeddings generated by the encoder, this module applies feature augmentation through random dropout (20%) to create distinct views. It then computes cosine similarities between positive pairs (same protein across views) and negative pairs (different proteins) (Eqs. 13-14). It optimizes the embeddings via a contrastive loss function (Eq.15). This process enhances the discrimination between interacting and non-interacting protein pairs.
  - Multi-Scale Graph Wavelet Transform (Section 2.2.2): This module extracts localized and global structural interaction patterns. It constructs a symmetric normalized matrix from the adjacency matrix (Eqs. 7-9), computes diffusion operators across biologically meaningful scales (step sizes k=1,2,3,4) (Eq. 10), and derives wavelet coefficients at these scales (Eqs. 11-12). The resulting multi-resolution feature representation captures information ranging from immediate binding interfaces (local) to pathway-level interactions (global).
b. Multi-view contrastive learning module: As described in (a), this module processes the hyperbolic node embeddings through feature augmentation with 20% random node dropout. By computing cosine similarities between positive pairs (same protein across views) and negative pairs (different proteins) following Eqs. 13-14, this module minimizes the contrastive loss function (Eq. 15) to generate discriminative embeddings that effectively separate interacting from non-interacting protein pairs.
c. Multi-scale graph wavelet transform: As described in (a), this module derives multi-resolution features through three computational stages:
  - Construction of a Symmetric Normalized matrix from the adjacency matrix (Eqs. 7-9).
  - Generation of multi-scale diffusion operators (Eq. 10) capturing neighborhood propagation dynamics across scales k=1,2,3,4.
  - Extraction of wavelet coefficients at the four discrete scales via spectral decomposition (Eqs. 11-12). The resulting feature representation encodes both immediate binding interfaces and pathway-level interaction patterns.

**Figure 1:**
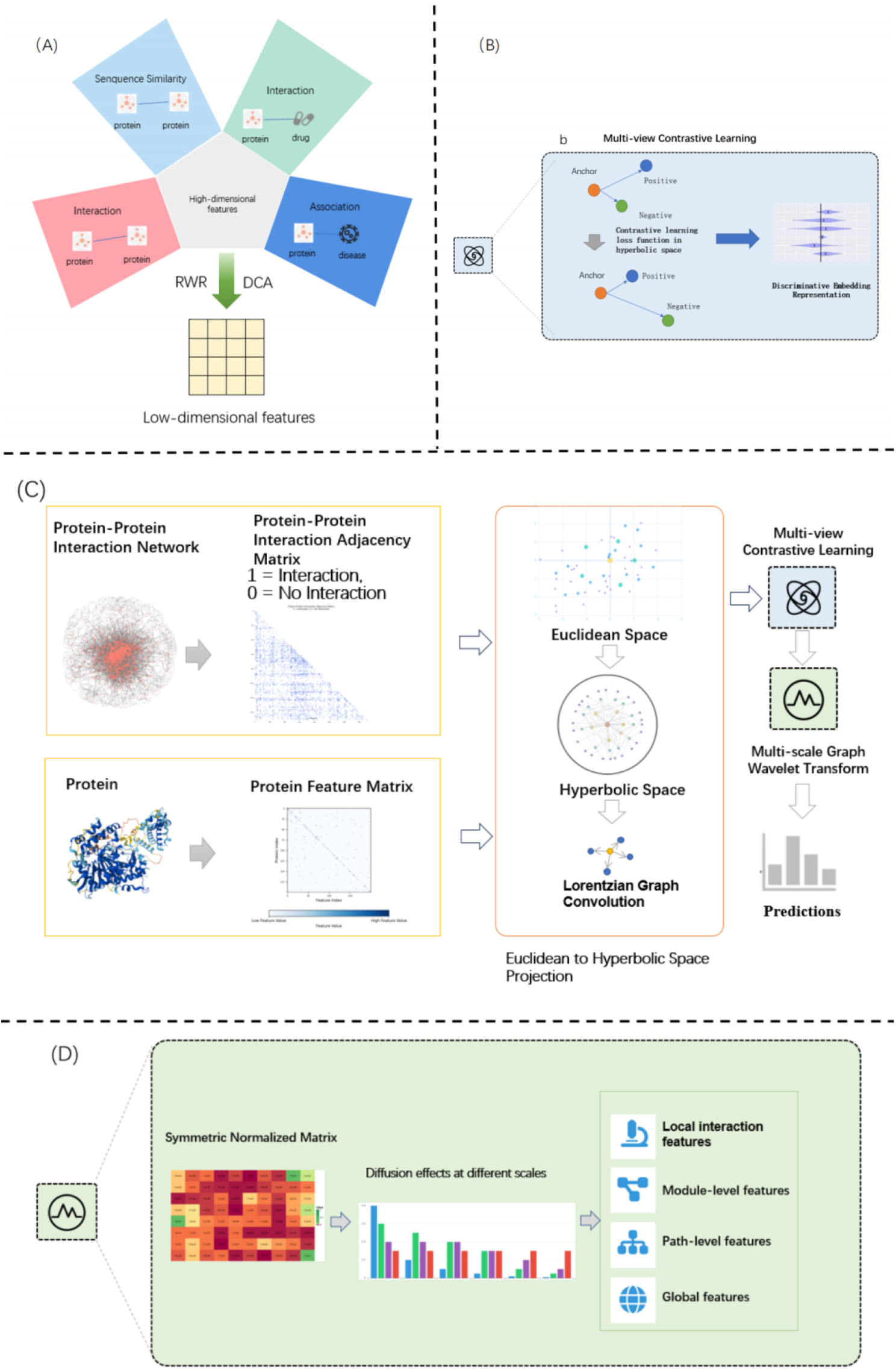
Architecture of HyWinNet for PPI prediction. A Biological multimodal diagram. Five modalities including sequence similarity, protein-protein interaction, protein-drug interaction and association, were used to generate unified protein representations. B Multi-view Contrastive Learning. This component employs an innovative dual-pathway architecture to learn robust protein representations. C Hyperbolic Graph Neural Network (Lorentz GCN). The geometric deep learning core operates in Lorentzian hyperbolic space with Manifold Projection, Hierarchical Propagation and Biological Advantages. D Multi-scale Wavelet Transform. This spectral analysis module extracts interaction patterns at four biologically meaningful scales.

#### 2.2.1. Lorentz space based graph neural network encoder(HyboNet)

Conventional Graph Neural Networks (GNNs) in Euclidean space exhibit limitations in capturing hierarchical relationships inherent in biological networks, such as scale-free topology and exponential growth of node neighborhoods[28]. To address this, we leverage **hyperbolic geometry**-specifically, **the Lorentz model**-due to its capacity to naturally embed hierarchical structures with minimal distortion. The Lorentz space cn is defined by the manifold:

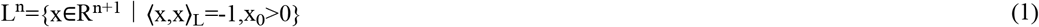

where the Lorentzian inner product is given by:

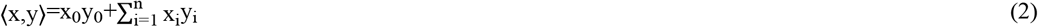

Here,**x**=(*x*_0_,*x*_1_,…,*x*_*n*_) denotes coordinates in the ambient space, with as the time-like dimension enforcing the manifold constraint.

##### Operational Framework

Given Euclidean input features for node, we first project them into via exponential mapping at a base point (the origin of hyperbolic space):

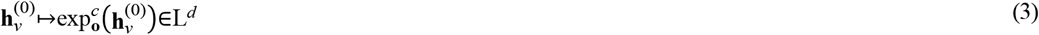

where c>0 is the curvature parameter controlling spatial distortion For each layerc>0, node embeddings are updated in three steps:

1. **Hyperbolic Neighborhood Aggregation**:Features from neighbors N(v) are aggregated using a permutation-invariant function:

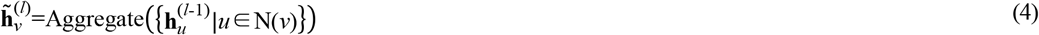 We use Aggregate, implemented as a weighted mean with attention coefficients.
2. **Lorentzian Feature Transformation**:The aggregated features are transformed via alinear operation in the tangent space at o:

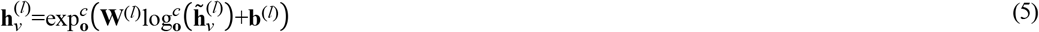 Where 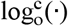 maps hyperbolic points to the tangent space, W^(l)^∈R^d’×d^ and b^(l)^∈R^d^’ are learnable parameters.

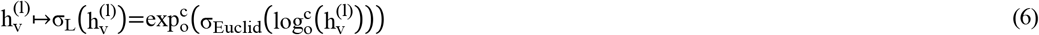

where Euclid is a standard activation (e.g., ReLU). The final output H^(L)^∈L^d^ provides hierarchical node embeddings that preserve the inherent tree-like structure of PPI networks.

#### 2.2.2. Multi-scale Spatial Wavelet Transformation via Symmetric Normalized Matrix

The traditional method is based on the graph Laplacian matrix L=D-A, whose spectral decomposition is L=UΛU^T^. Instead, multiscale neighborhood relationships are explicitly modeled by diffusion via symmetric normalized adjacency matrices to avoid the computational bottleneck of feature decomposition. The number of diffusion steps s directly corresponds to the spatial scale, ranging from local (s=1) to global (s=K), and naturally aligns with biological hierarchies (for example, protein domains composed of amino acid residues).

The first step is to build the symmetric normalized matrix. Given an undirected graph g=(v,ε) and an adjacency matrix A∈{0,1}^N×N^, construct the adjacency matrix with self-loop:

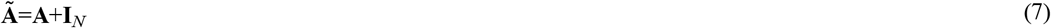

The degree matrixD∈R^N×N^ is defined as a diagonal matrix whose elements satisfy:

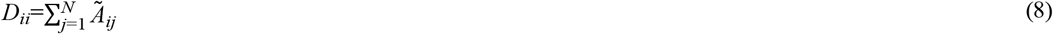

The symmetric normalized matrix P∈R^N×N^ is defined as:

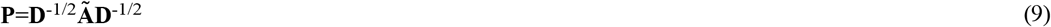

Subsequently, a multiscale diffusion operator is constructed using a symmetric normalized matrix and a predefined range of diffusion scales.

Given a set of scale parameters S={s_1_,s_2_,…,s_K_}, the scale-dependent diffusion operator T_s_∈ R^N×N^ is formulated by raising the symmetric normalized matrix to the power s:

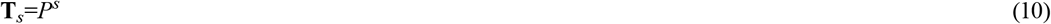

where s∈S controls the number of diffusion steps and larger s captures more global neighborhood information.

Finally, the generated wavelet coefficients are used with the feature matrix for graph wavelet feature extraction. Given the node feature matrix X∈R^N×d^, the multiscale graph wavelet feature Z ∈R^N×(K.d)^ is defined as:

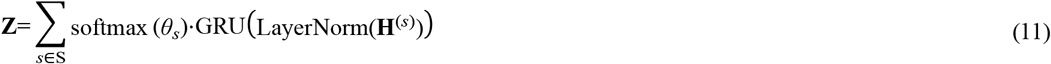

- S denotes the pre-defined set of diffusion scales (e.g., S={1,2,4,8})
- θ_s_∈R represents the learnable importance weight for scale s
- softmax 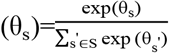 normalizes scale weights
- H^(s)^=P^s^X contains the diffusion features at scale s
- LayerNorm(·) applies feature-wise normalization
- GRU(·) denotes the Gated Recurrent Unit that models cross-scale interactions

#### 2.2.3. Contrastive Learning Module

In the contrastive learning module, we generate an augmented view G^’^=(X^’^,A^’^) by independently applying a 20% random dropout to the feature matrix and the adjacency matrix, thus introducing perturbations that enhance the robustness of representation learning. The original view G=(X,A) is fed into the model to generate two node embeddings Z^’^ and Z, which are subsequently used to compute the alignment 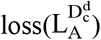 (Eqation [12]) and outer shell isotropy 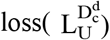 (Eqation [15]). The notation 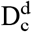 denotes the d-dimensional Poincaré ball model, where N represents the number of nodes. The core component 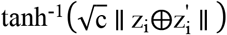 computes the hyperbolic distance in the Poincaré ball model, where:

- ⨁ denotes Möbius addition, which is the vector addition operation in hyperbolic space
- c>0 is the curvature parameter of the Poincaré ball
- ||.|| represents the Euclidean norm

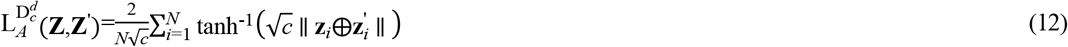

Equation(13) computes the tangent space statistics for subsequent KL divergence loss, where denotes the hyperbolic logarithm map with the origin as base point, and 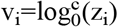.

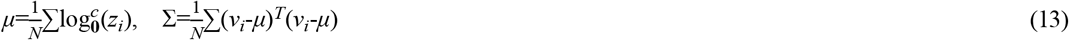

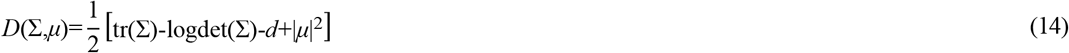

where:

- μ: The mean vector of the features projected onto the tangent space, as defined in Equation [13].
- Σ: The covariance matrix of the tangent space features, also computed according to Equation [13].
- d: The dimensionality of the embedding space.
- tr(): The trace operator, which computes the sum of the diagonal elements of a matrix.
- logdet(): The logarithm of the determinant of the covariance matrix

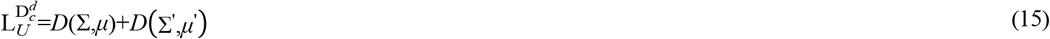

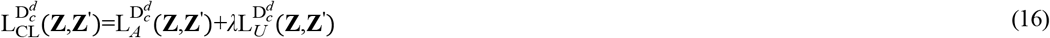

λ is a tunable scaling parameter.

### 2.3. Protein-Protein Interaction Prediction

The feature representations produced by the encoder are employed to compute the squared Lorentzian distance between pairs of nodes in hyperbolic space, which is subsequently used as the node interaction score.

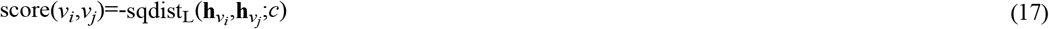

where 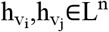 are the embeddings of nodes v_i_ and v_j_ in Lorentz hyperbolic space, sqdist_L_(.) is the squared distance function in Lorentz hyperbolic space, and c>0 is the curvature parameter of Lorentz space.

## 3. Experiments and Results

Since protein-protein interaction (PPI) prediction can be formulated as a binary classification task, we evaluate model performance using two key metrics:

- ROC-AUC (Receiver Operating Characteristic - Area Under the Curve): This fundamental evaluation metric for binary classifiers measures the model’s ability to correctly rank positive and negative samples. The ROC curve is generated by plotting the True Positive Rate (TPR) against the False Positive Rate (FPR) at various classification thresholds, with the Area Under this Curve (AUC) providing a scalar value representing overall performance (hereafter referred to as AUC). An AUC of 1.0 indicates perfect classification, while 0.5 represents random guessing.

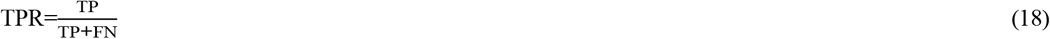
  - TP (True Positives): Number of positive instances correctly classified
  - FN (False Negatives): Number of positive instances incorrectly classified as negative

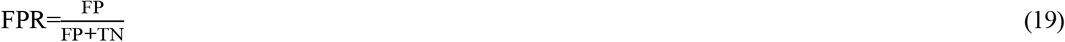
  - FP (False Positives): Number of negative instances incorrectly classified as positive
  - TN (True Negatives): Number of negative instances correctly classified

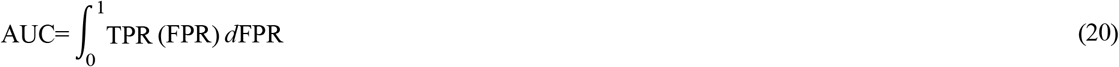
- Average Precision (AP) is a widely adopted evaluation metric in information retrieval and machine learning, especially for class-imbalanced binary classification problems[30]. It quantifies the ranking performance of positive-class instances by averaging precision values at multiple recall levels, commonly abbreviated as AP in subsequent discussions.

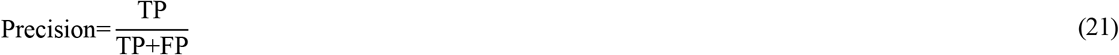

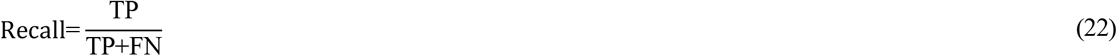

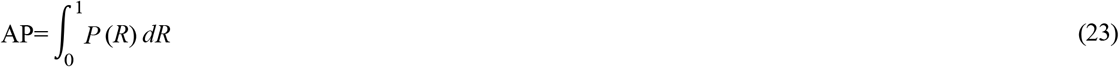
  - P(R): Precision as a function of Recall
  - integration interval R∈[0,1]

In our protein-protein interaction (PPI) prediction study, we constructed the training dataset using experimentally confirmed PPIs as positive samples and randomly paired non-interacting proteins as negative samples. The HyWinNet model was implemented using Luo et al.’s benchmark dataset, incorporating a multi-scale wavelet transform with four distinct scales (1, 2, 3, and 4) for feature extraction. For the contrastive learning framework, we generated comparative views by randomly dropping 20% of features and setting the temperature parameter to 0.2. The model was trained using the Adam optimizer with a learning rate of 0.001 and an early stopping mechanism, running for up to 2000 epochs to ensure convergence while preventing overfitting.

### 3.1. Benchmark Comparison

We compared our model with five state-of-the-art baseline methods, as summarized in Table 1. To ensure a fair comparison, all models were evaluated on the same dataset. Our method consistently outperformed all five baselines across the evaluated metrics:

**Table 1.**
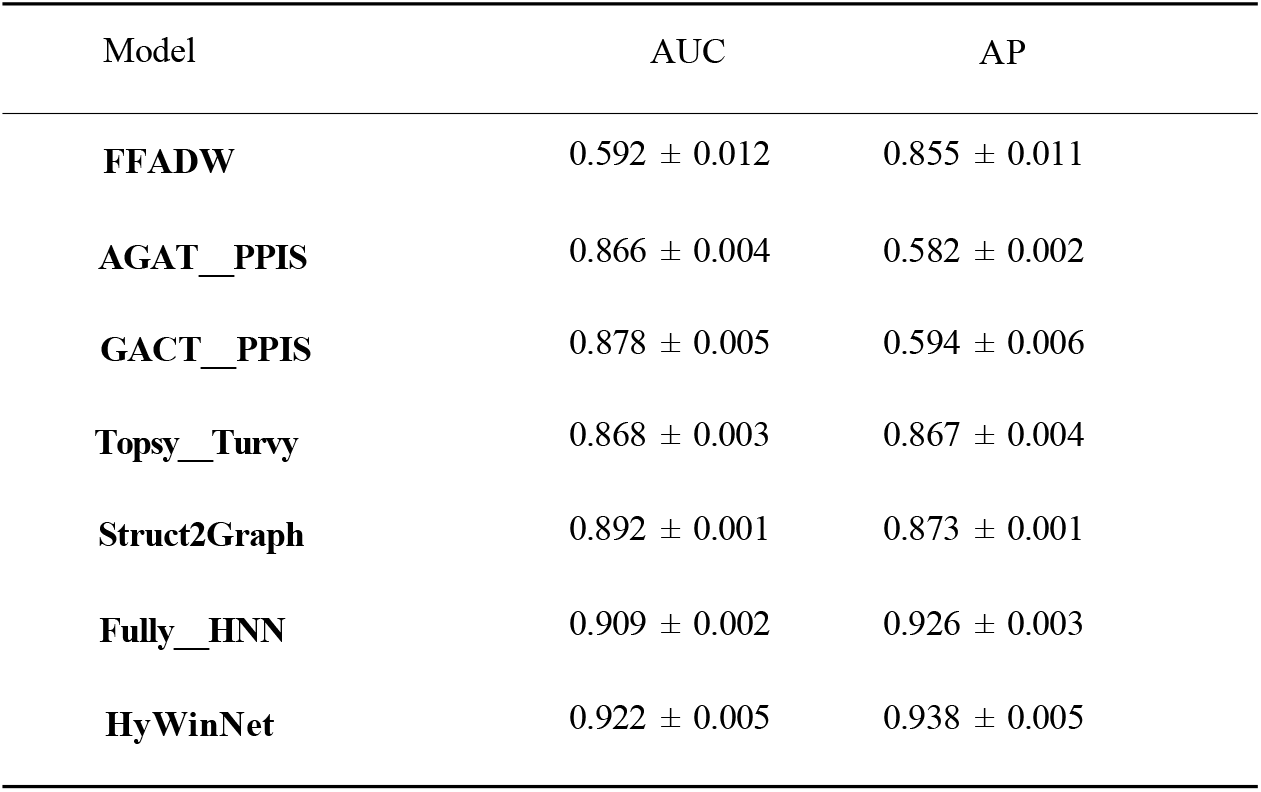
Comparison of performance with benchmark models.

- **FFADW** employs a weighted fusion strategy that integrates sequence similarity, quantified by the Levenshtein distance, and network similarity, computed via a Gaussian kernel function. The fused similarity matrix is then processed by the Attributed DeepWalk algorithm, which incorporates both topological and attribute information to generate robust low-dimensional embeddings for protein-protein interaction prediction[31].
- **AGAT_PPIS** is an augmented graph attention network-based predictor for protein-protein interaction sites, incorporating initial residual and identity mapping[32].
- **GACT_PPIS** was designed to predict protein-protein interaction sites by leveraging combined information from both protein sequences and structures[33].
- **Struct2Graph** is a structure-based prediction method that converts 3D protein structures into graphs and applies graph-based learning to infer interactions[34].
- **Topsy_Turvy** is a sequence-based approach that utilizes amino acid sequences and evolutionary information for interaction prediction, enhanced with transfer learning techniques to improve performance on sequence data[35].
- **Fully_HNN** is a fully hyperbolic framework based on the Lorentz model, which formalizes fundamental neural network operations through Lorentz transformations (including boost and rotation) to construct hyperbolic networks[28].

As shown in Table 1, the proposed model achieved an AUC of 0.922 and an AP of 0.938, outperforming all competing state-of-the-art methods. Specifically, it yielded AUC improvements ranging from 0.013 to 0.056 and AP gains between 0.012 and 0.356. The superior performance of HyWinNet can be attributed to three key innovations:

(1) projecting features into hyperbolic space under the Lorentz model to better capture their intrinsic hierarchical structure, (2) employing multi-scale wavelet transforms on the adjacency matrix to simultaneously extract both local and global structural patterns, and (3) utilizing contrastive learning with multiple views to effectively separate positive and negative samples in the embedding space.

As shown in Fig. 2, the model maintains robust operational characteristics beyond aggregate metrics. The ROC curve (Fig. 2a, AUC=0.922) and precision-recall curve (Fig. 2b, AUPR=0.938) collectively demonstrate:

**Figure 2:**
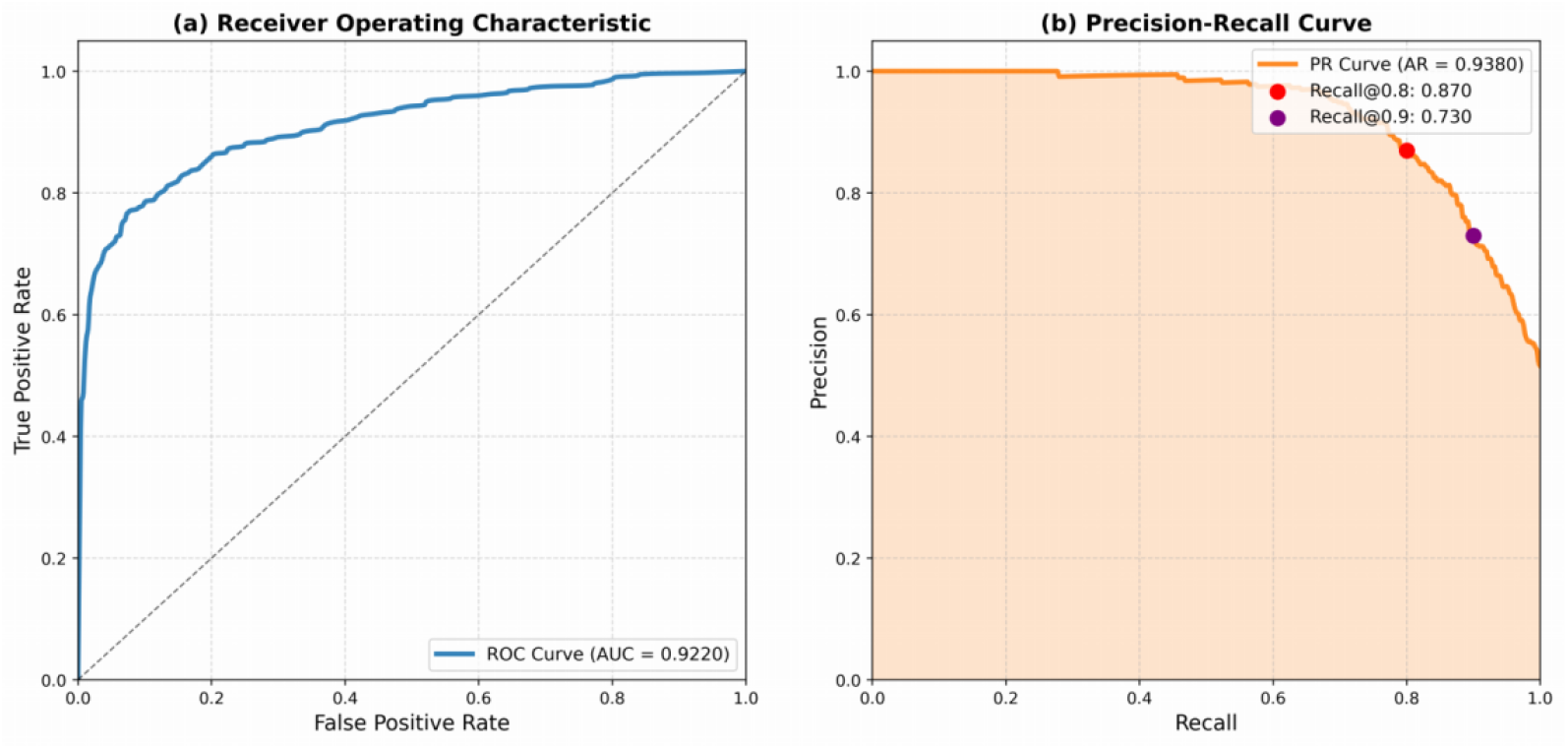
Model performance evaluation. (a) Receiver Operating Characteristic (ROC) curve demonstrating strong discriminative ability (AUC = 0.9220, 95% CI: 0.915-0.929), significantly outperforming random guessing (dashed line). (b) Precision-Recall curve showing robust performance across recall levels (Average Precision = 0.9380), with precision values of 0.870 and 0.730 at recall levels of 0.8 and 0.9, respectively. Both curves were smoothed using PCHIP interpolation, with shaded regions indicating ±1 standard deviation from 1,000 bootstrap samples. The model maintains high precision even at stringent recall thresholds.

- The model maintains diagnostic precision across the spectrum of sensitivity requirements.
- Optimal operating points cluster at higher specificity levels.
- Performance degradation occurs gradually rather than precipitously at extreme recall levels.

### 3.2. Ablation experiments

#### 3.2.1. Ablation experiment on the encoder

We conducted ablation studies on both datasets, first targeting the Lorentz space-based graph neural network encoder. Under the experimental configuration with the “luo” dataset, curvature scales [1, 2, 3, 4], and temperature parameter 0.2, we systematically replaced the encoder with three variants:

- Euclidean-based GCN
- Hyperboloid-based HGCN
- Lorentz space-based Fully

As demonstrated in Fig. 3, both AUC and AP metrics exhibited consistent degradation across all encoder replacements. Specifically:

**Figure 3:**
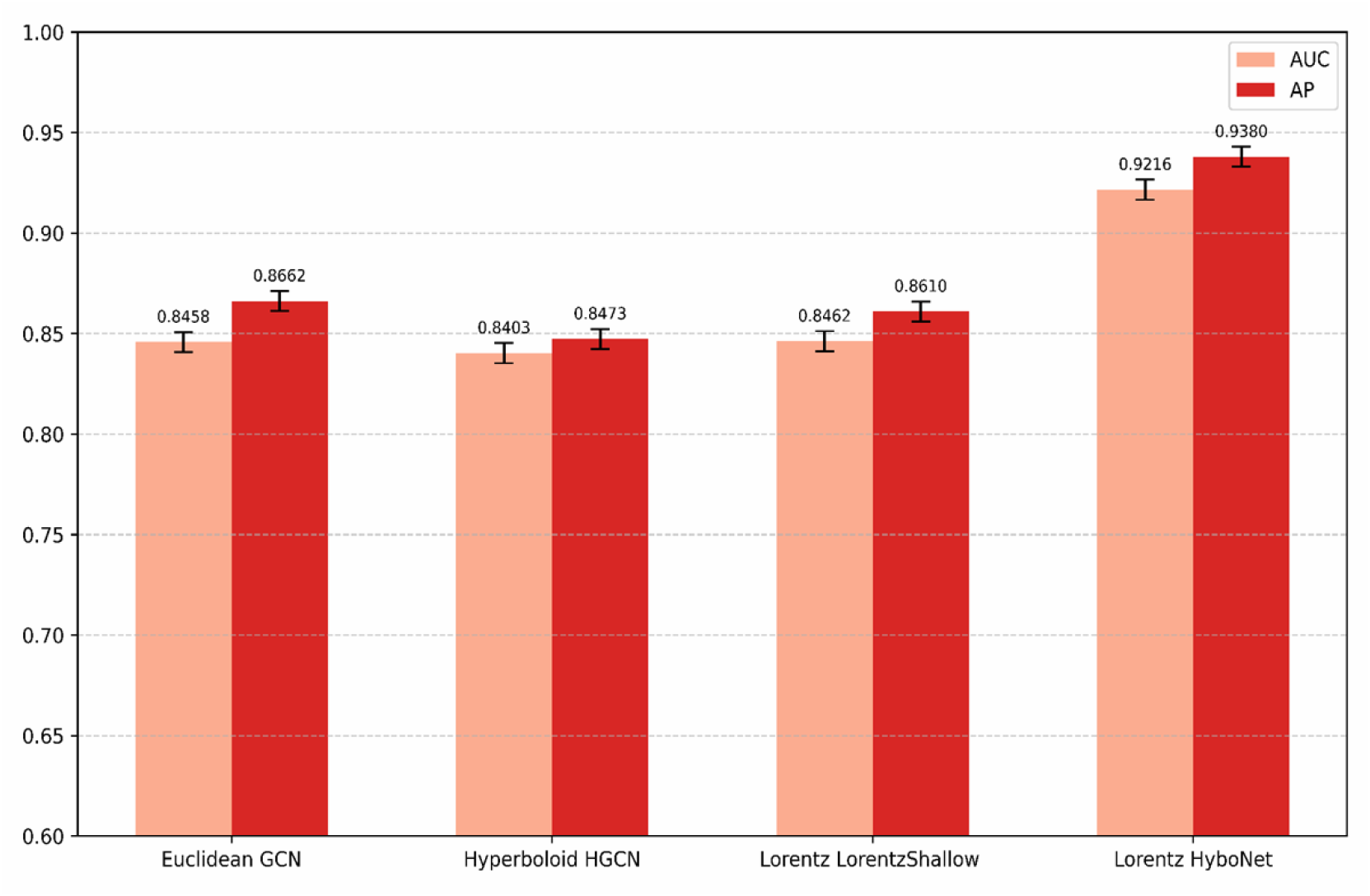
Comparative Analysis of AUC and AP Evaluation Metrics Across Diverse Encoders on the Luo Dataset evaluation metrics of the GWT module are also higher than those without the GWT module. We believe that in the Zeng dataset, the features of proteins combine the relationships of biological networks. This causes the GWT to capture the local and global information of the graph too deeply and lose some superficial information, ultimately affecting the evaluation metrics.Overall, it can be inferred that the GWT module is capable of improving the predictive performance of the model.

- AUC: Maximum difference 0.0817, minimum difference 0.0758
- AP: Maximum difference 0.0907, minimum difference 0.071

For the ablation of the graph neural network encoder based on Lorentz space, with scale [1, 2, 3, 4] and temperature parameter 0.2, we changed the encoder toGCN under Euclidean, HGCN under Hyperboloid, and LorentzShallow under Lorentz for the experiments as shown in Fig. 3, and after the The values ofAUC and AP are decreased after changing the encoder.

We conducted experiments on the “Zeng” dataset with curvature scales [1, 2, 3, 4] and temperature parameter 0.2, systematically replacing the encoder with three implementations:

- Euclidean-based GCN
- Hyperboloid-based HGCN
- Lorentz space-based LorentzShallow

As shown in Fig. 4, while AUC and AP metrics demonstrated improvements under the LorentzShallow encoder, both metrics exhibited degradation with other encoders. Specifically:

**Figure 4:**
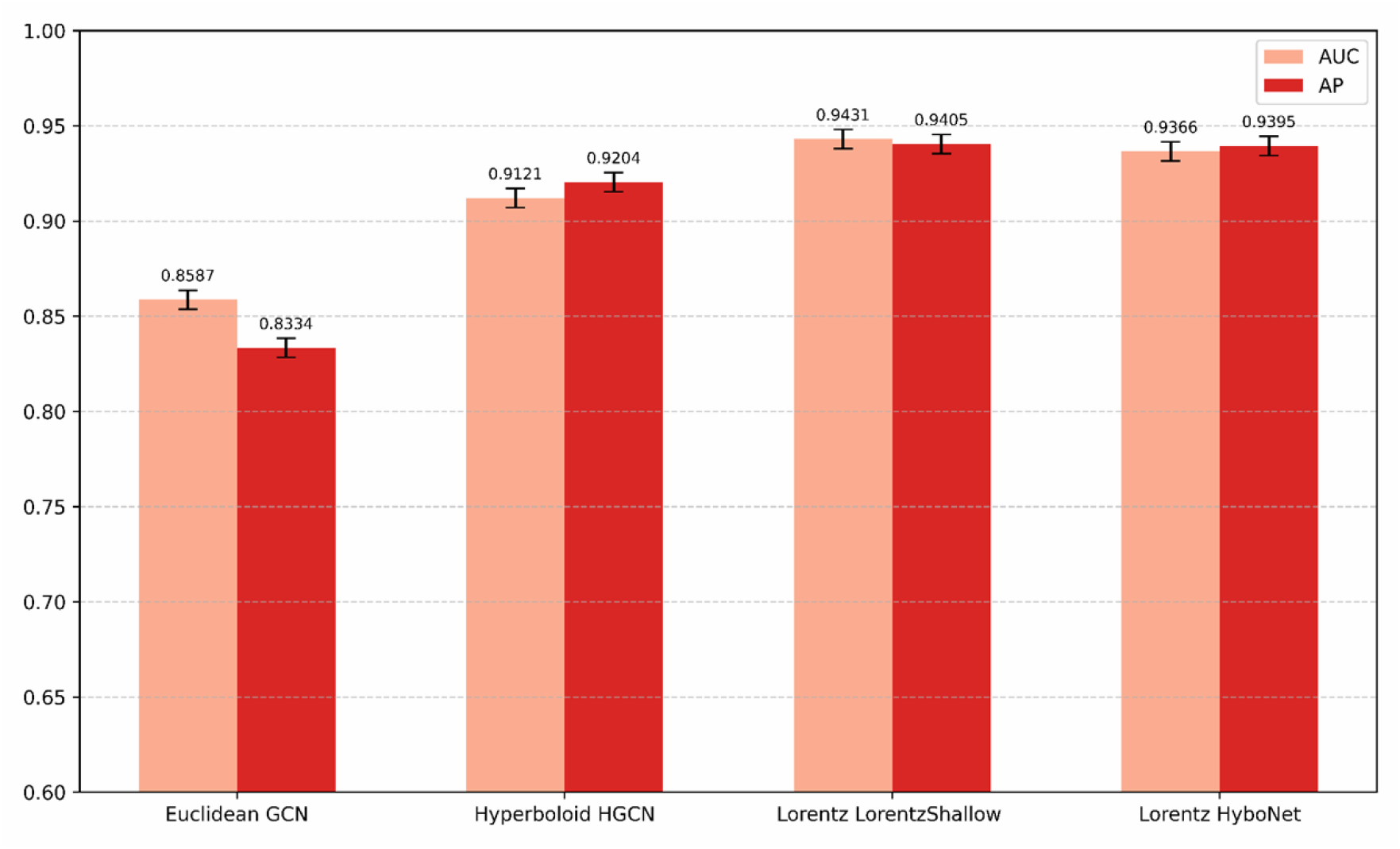
Comparative Analysis of AUC and AP Evaluation Metrics Across Diverse Encoders on the Zeng Datase

- AUC: Maximum difference 0.0799, minimum difference 0.0245
- AP: Maximum difference 0.1061, minimum difference 0.0191

This performance enhancement with LorentzShallow suggests the pre-existing structural features in protein characterization within the “Zeng” dataset inherently align with Lorentzian geometric representations, creating synergistic effects unavailable in Euclidean/Hyperbolic paradigms.

For robustness assessment, we performed bootstrap analysis with 1,000 resamples to estimate 95% confidence intervals for both AUC and AP metrics, with results visualized in Fig.5. The LorentzShallow encoder demonstrated the narrowest confidence intervals for AUC scores, while HyboNet achieved the most precise estimation (smallest CI range) for AP metrics. Notably, HyboNet’s AUC confidence interval spanned merely 0.31%, further evidencing its exceptional robustness as an encoder architecture.

**Figure 5:**
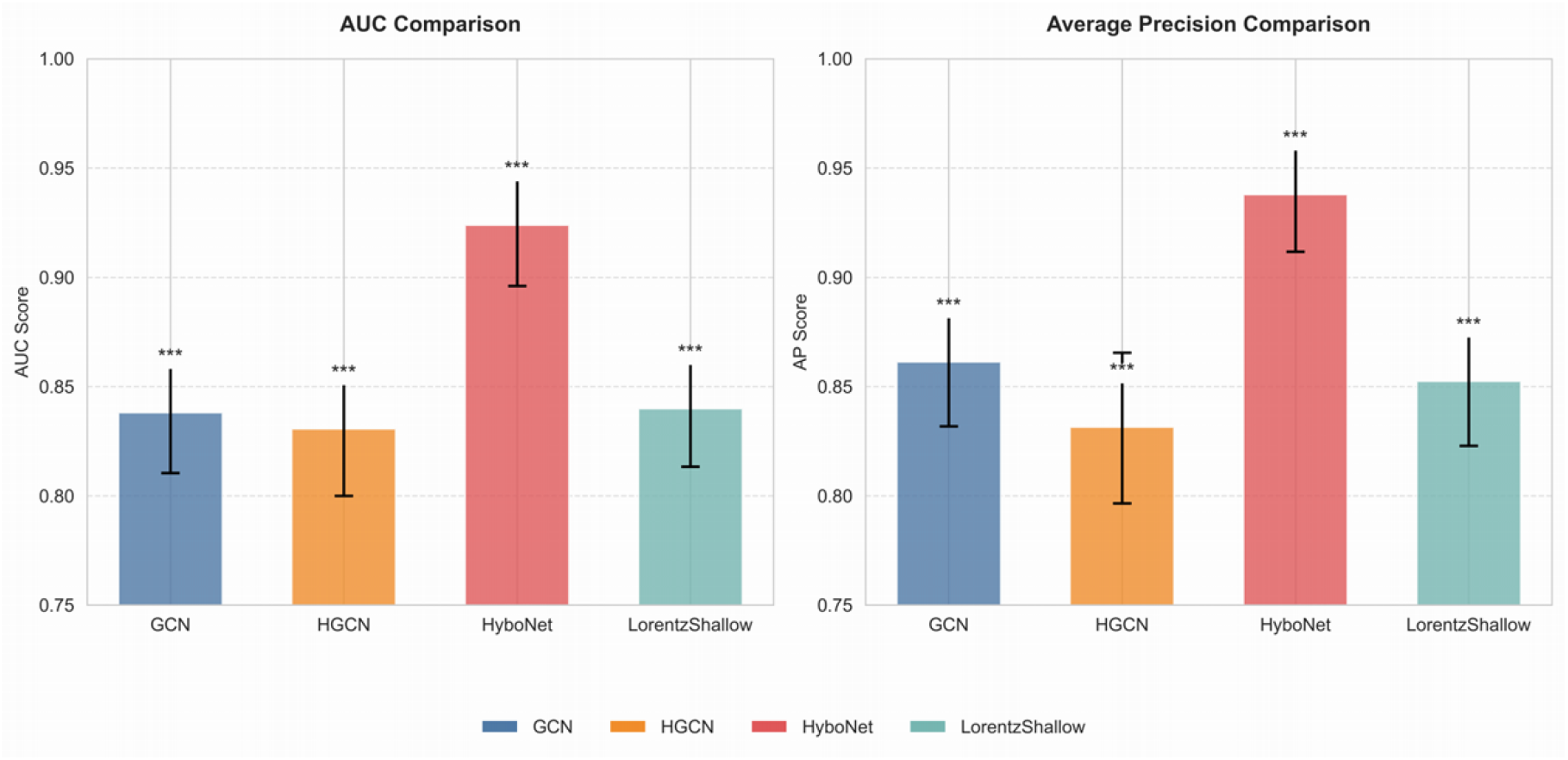
Robustness Assessment via Bootstrap Resampling: 95% Confidence Intervals for AUC and AP Metrics.

#### 3.2.2. Ablation experiment on the GWT module

This section will conduct ablation experiments on the GWT module under different datasets and encoders, as shown in Table 2. Except for the LorentzShallow encoder, the evaluation metrics of the GWT module are higher than those without the GWT module in the Zeng dataset and the same encoder. In the Luo dataset and the same encoder, the evaluation metrics of the GWT module are also higher than those without the GWT module. We believe that in the Zeng dataset, the features of proteins combine the relationships of biological networks. This causes the GWT to capture the local and global information of the graph too deeply and lose some superficial information, ultimately affecting the evaluation metrics.Overall, it can be inferred that the GWT module is capable of improving the predictive performance of the model.

**Table 2.** GWT ablation experiment.

| Dataset | Encoder | Module | AUC | AP |
| --- | --- | --- | --- | --- |
| **Zeng** | GCN | GWT+CL | 0.8587 | 0.8334 |
| **Zeng** | GCN | CL | 0.7995 | 0.7854 |
| **Zeng** | HGCN | GWT+CL | 0.9121 | 0.9204 |
| **Zeng** | HGCN | CL | 0.8955 | 0.9009 |
| **Zeng** | LorentzShallow | GWT+CL | 0.9431 | 0.9405 |
| **Zeng** | LorentzShallow | CL | 0.9527 | 0.9518 |
| **Zeng** | HyboNet | GWT+CL | 0.9366 | 0.9395 |
| **Zeng** | HyboNet | CL | 0.9208 | 0.9190 |
| **Luo** | GCN | GWT+CL | 0.8458 | 0.8662 |
| **Luo** | GCN | CL | 0.7736 | 0.7831 |
| **Luo** | HGCN | GWT+CL | 0.8403 | 0.8473 |
| **Luo** | HGCN | CL | 0.7752 | 0.7997 |
| **Luo** | LorentzShallow | GWT+CL | 0.8462 | 0.8610 |
| **Luo** | LorentzShallow | CL | 0.8402 | 0.8549 |
| **Luo** | HyboNet | GWT+CL | 0.9216 | 0.9380 |
| **Luo** | HyboNet | CL | 0.9151 | 0.9309 |

#### 3.2.2. Ablation experiment on the GWT module

This section will conduct ablation experiments on the GWT module under different datasets and encoders, as shown in Table 2. Except for the LorentzShallow encoder, the evaluation metrics of the GWT module are higher than those without the GWT module in the Zeng dataset and the same encoder. In the Luo dataset and the same encoder, the evaluation metrics of the GWT module are also higher than those without the GWT module. We believe that in the Zeng dataset, the features of proteins combine the relationships of biological networks. This causes the GWT to capture the local and global information of the graph too deeply and lose some superficial information, ultimately affecting the evaluation metrics.Overall, it can be inferred that the GWT module is capable of improving the predictive performance of the model.

#### 3.2.3. Ablation experiments on contrastive learning

This section conducts an ablation study on the contrastive learning module using the Zeng dataset, as depicted in Fig. 6 and Fig. 7. The results demonstrate that, under the same encoder configuration, the model incorporating the contrastive learning module achieves consistently superior evaluation metrics compared to the baseline without the module, with observed maximum and minimum performance differences of [insert values]. These findings suggest that the contrastive learning module significantly enhances the model’s predictive capabilities.

**Figure 6:**
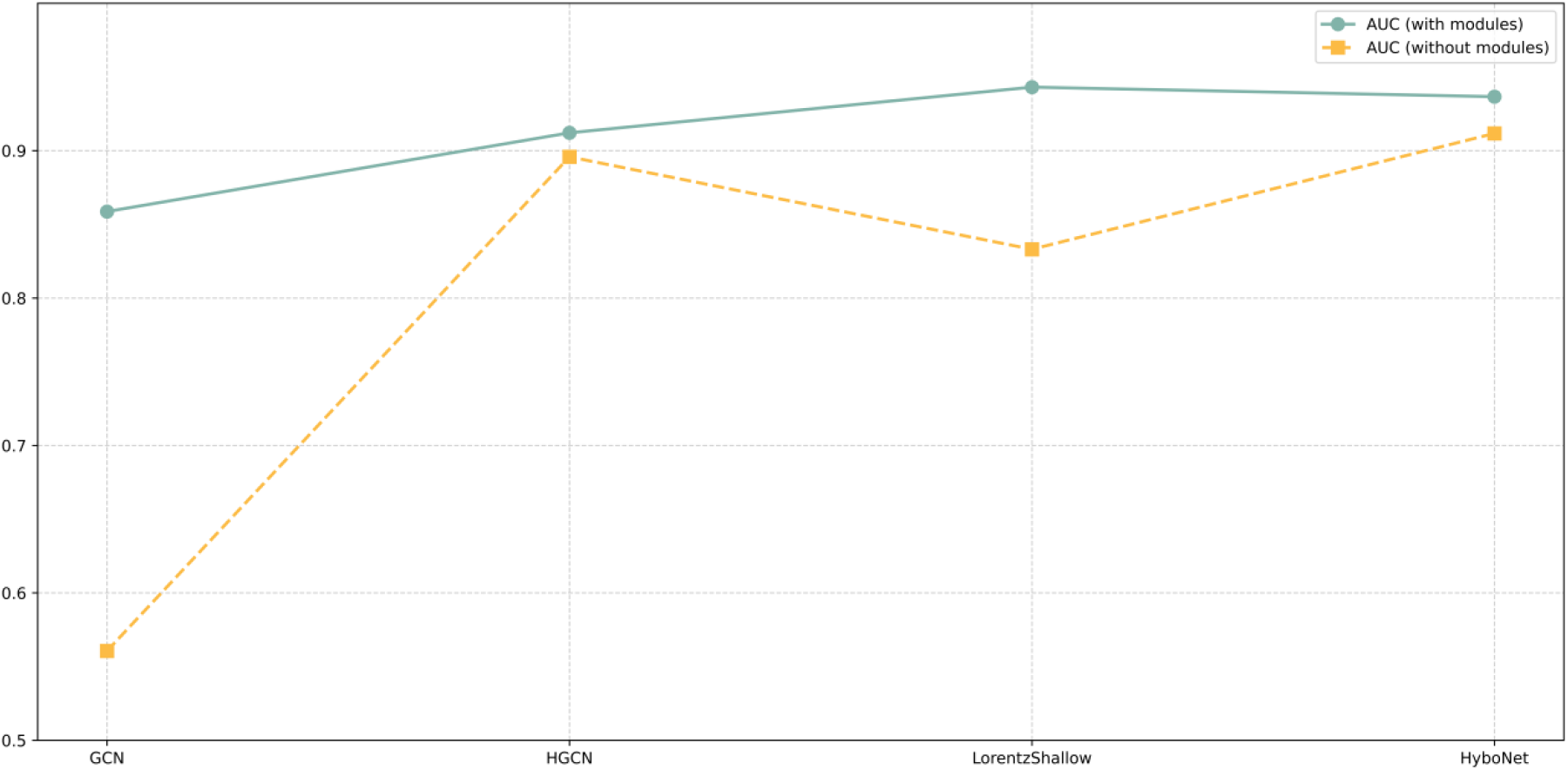
The Effect of Contrastive Learning Module Removal on Model Performance (AUC Scores)

**Figure 7:**
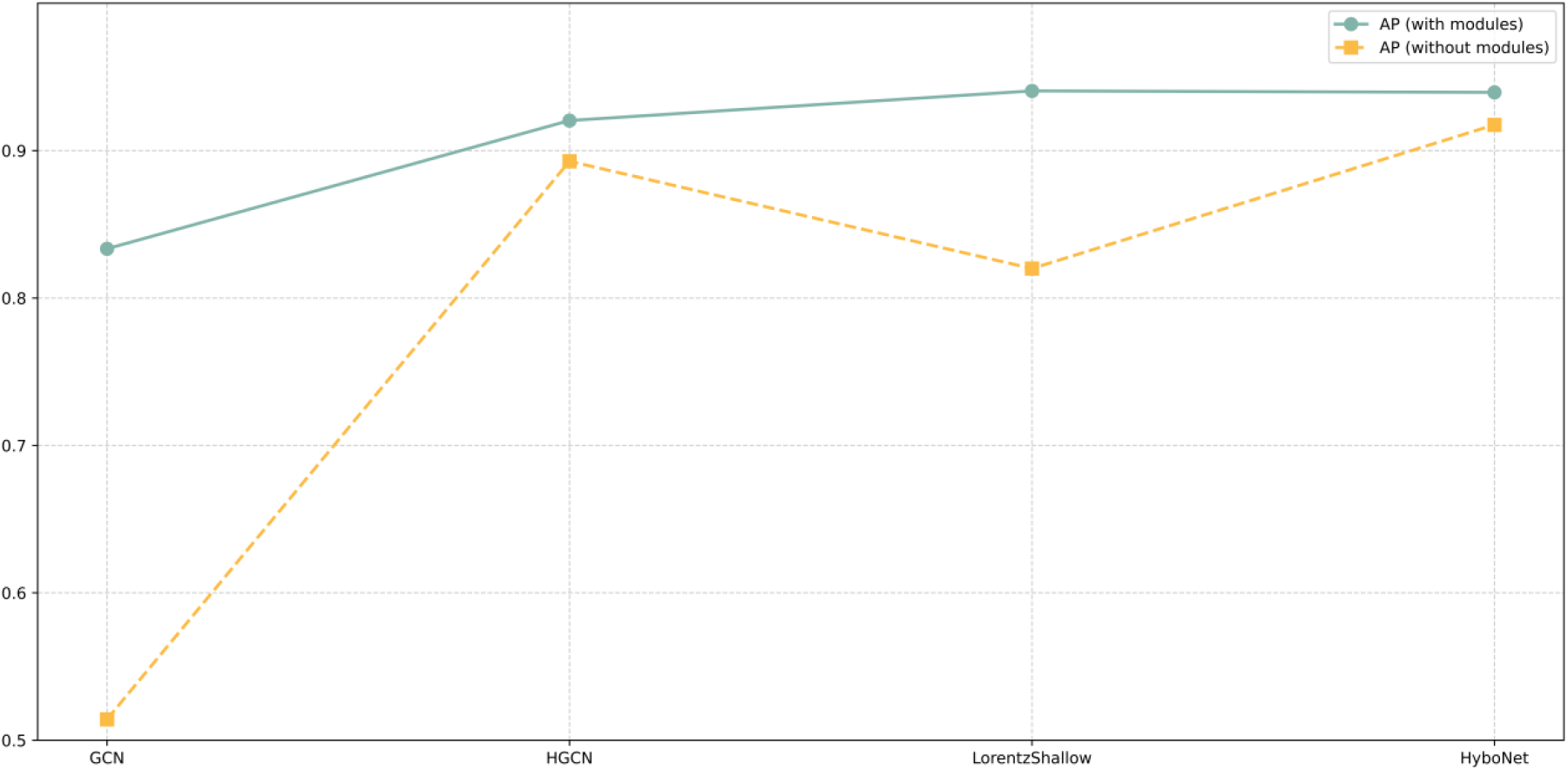
The Effect of Contrastive Learning Module Removal on Model Performance (AP Scores)

### 3.3. Hyperparameter Sensitivity Analysis

#### 3.3.1. Scale parameter sensitivity in wavelet transform analysis

We have performed a sensitivity analysis of hyperparameters of We conducted a sensitivity analysis of hyper-parameters in HyWinNet to identify optimal predictive performance. First, we systematically examined the number of scales and their corresponding values in the graph wavelet transform, restricting our exploration to 2-4 scales with individual scale values not exceeding 7. As illustrated in Fig.8, increasing the number of scales from 2 to 3 induced a substantial performance enhancement, suggesting that configurations with 3-4 scales capture broader and deeper structural patterns. When testing a 4-scale configuration with values (1, 2, 3, 4), both AUC and AP stabilized, outperforming smaller-scale variants. This optimal configuration demonstrates HyWinNet’s enhanced capacity to model hierarchical topological relationships among biomolecules.

**Figure 8:**
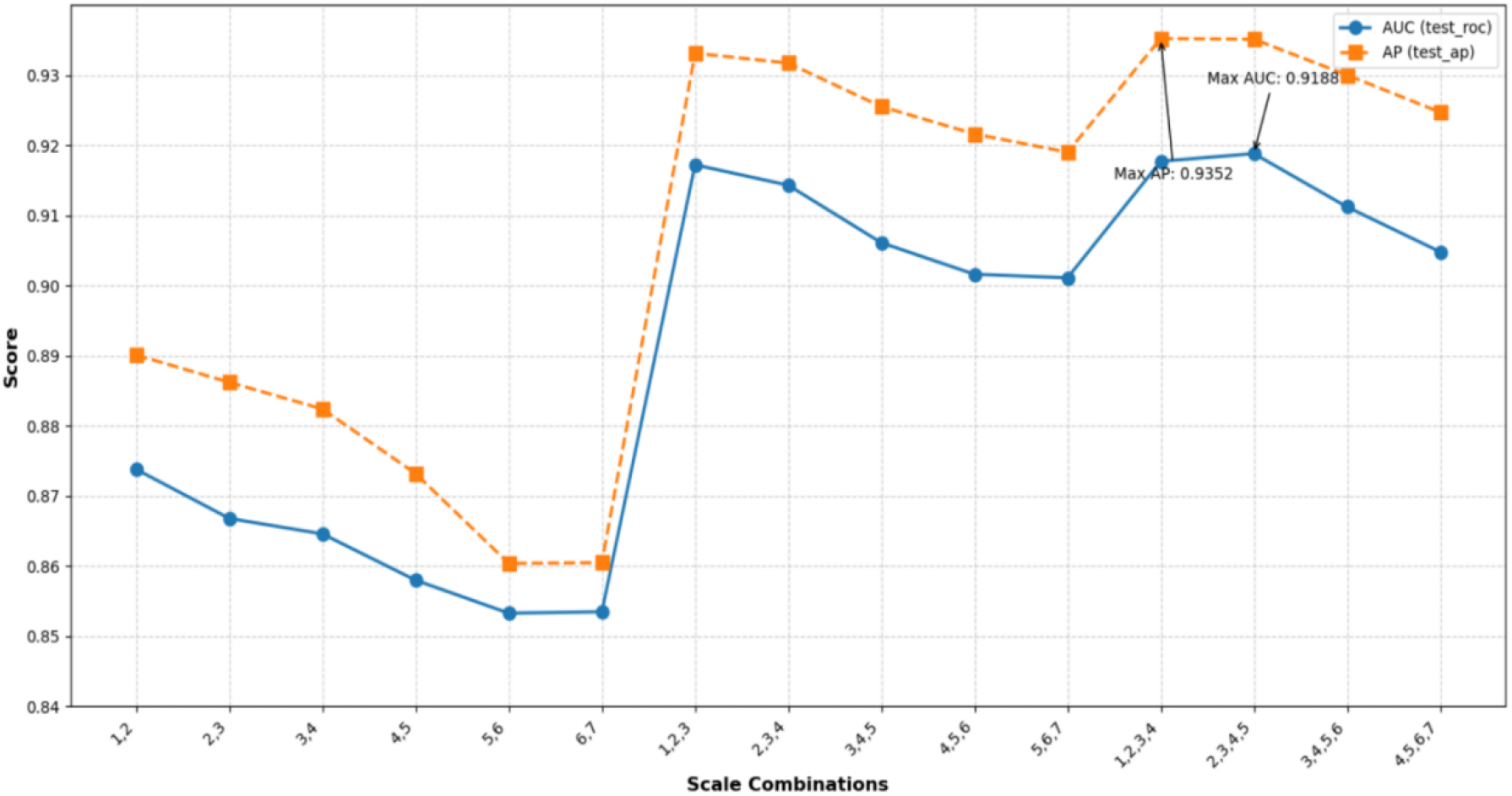
Values of AUC and AP under different scale lists

#### 3.3.2. Sensitivity Analysis of Data Splitting Ratio Parameters

To evaluate the impact of the training set ratio on model performance, we designed a systematic controlled experiment: adjusting the training set ratio (50%-90%) while correspondingly modifying the validation set ratio. The experiment compared two data processing strategies: balanced data (achieved through undersampling for class balance) and imbalanced data (retaining the original distribution). This dual-metric (AUC/AP) and dual-strategy (balanced/imbalanced) cross-validation was visually represented through radar chart polygon areas and shaded regions to illustrate stability. As shown in Fig. 9, performance declined significantly when the training ratio was <75%. Peak performance (AUC=0.921, AP=0.934) was achieved at 80%-85% training ratios. When exceeding 85%, insufficient test set size led to performance degradation. Notably, the imbalanced data strategy consistently outperformed the balanced strategy across all partitioning ratios, likely because the original data distribution reflects real-world scenarios and preserves the inherent data structure. Furthermore, the highly consistent trends between AUC and AP metrics validated the model’s robustness.

**Figure 9:**
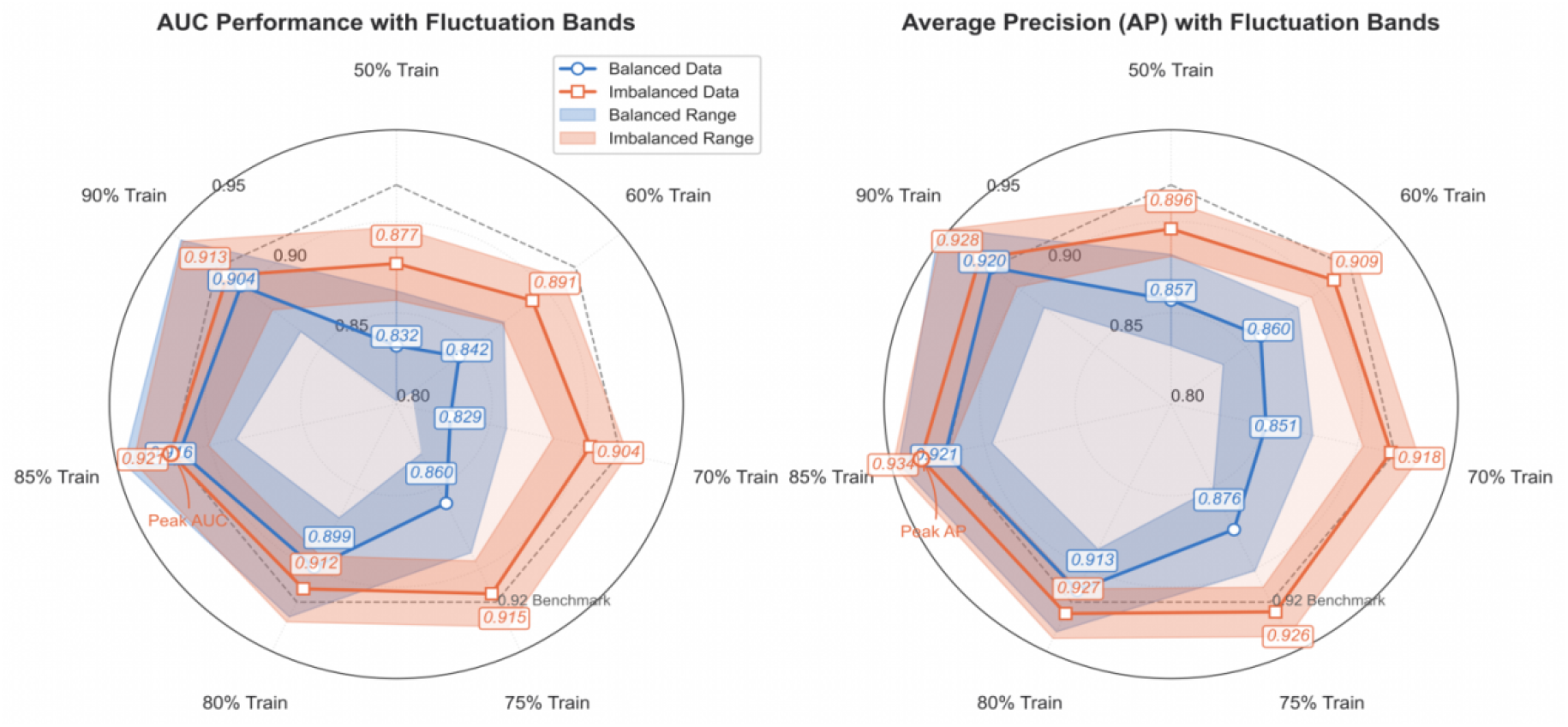
Radar Charts Comparing Model Performance: Imbalanced vs. Balanced Data with Fluctuation Bands

#### 3.3.3. Sensitivity Analysis of the Temperature Parameter in Contrastive Learning

Fig.10 reveals the significant sensitivity of model performance to the temperature parameter (τ) in the contrastive learning framework. Key observations include:

- The optimal operating point occurs at τ=0.05, achieving peak performance on both validation and test sets (ROC AUC= 0.944, AP= 0.953).
- A secondary performance peak emerges at τ=0.45 (Test AP= 0.947), suggesting the existence of potentially complementary optimization regions.
- The interval τ∈[0.4,1.0] demonstrates high parameter sensitivity, where minor adjustments (±0. 1) yield performance fluctuations up to 0.03. This non-linear response highlights the critical importance of precise temperature tuning.

**Figure 10:**
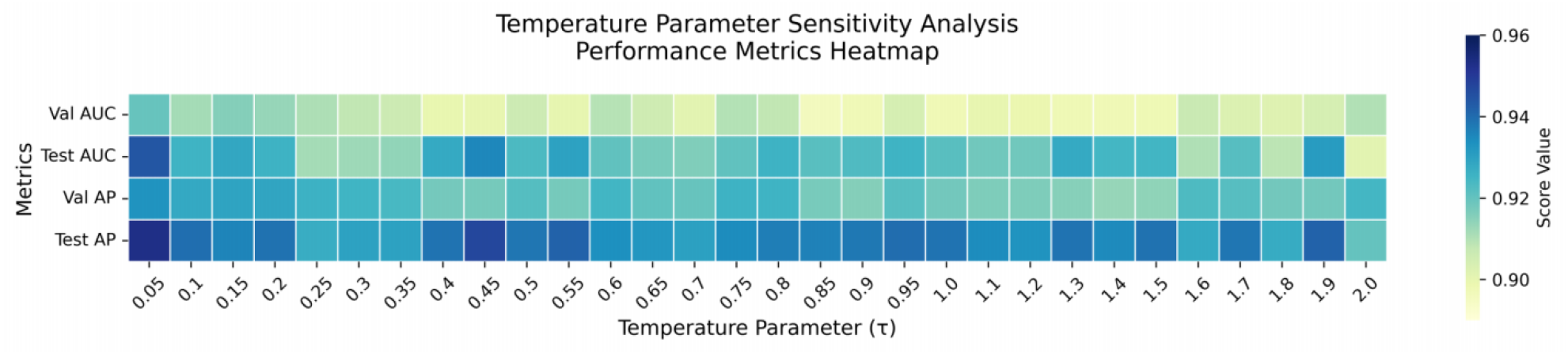
Temperature Parameter Sensitivity Analysis

Validation and test metrics exhibit congruent response patterns, confirming the reliability of observed effects. The consistent superiority of lower temperatures (τ<0.5) supports theoretical expectations that temperature scaling enhances feature discrimination in contrastive frameworks.

Based on these findings, we recommend τ=0.05 as the default configuration, with τ=0.45 as a viable alternative for specific application scenarios.

### 3.4. Statistical Significance Validation via Permutation Tests

To verify whether the protein interaction patterns captured by HyWinNet surpass random noise, we employ permutation tests to evaluate the statistical significance of the model, as shown in Fig.11 The specific design is as follows:

**Figure 11:**
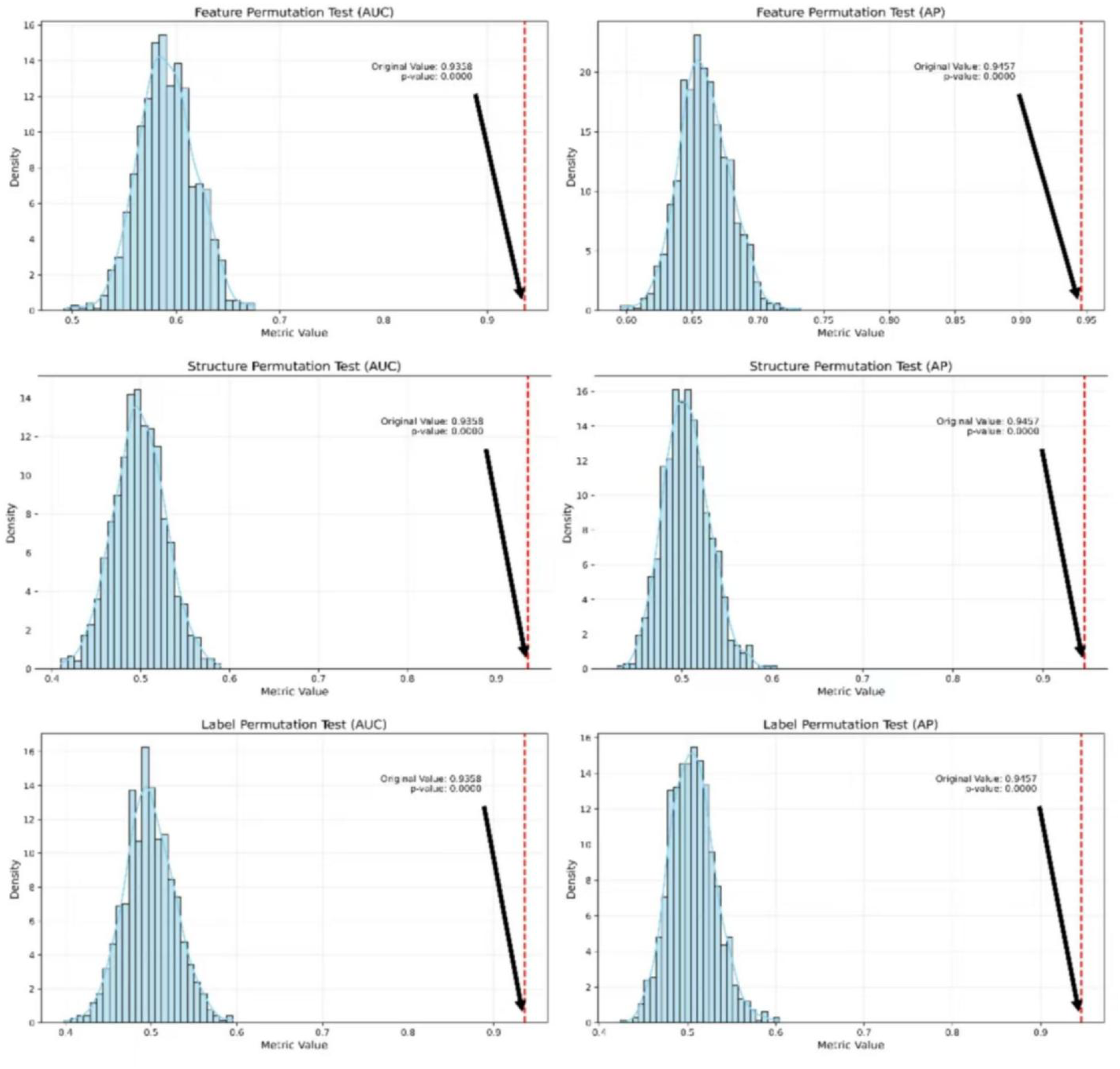
The permutation results illustrate the effects of feature shuffling, network rewiring, and label randomization on model performance (quantified by AUC and AP metrics).

#### Feature permutation

Randomly shuffle the rows of the node feature matrix X (i.e., protein features), disrupting the association between features and structure.

#### Structure permutation

Randomly rewire the adjacency matrix A (while preserving the degree distribution), destroying the true topological relationships.

#### Label permutation

Randomly flip PPI labels (positive/negative samples), breaking the causal relationship between labels and data.

#### Test metric

Compute the average performance decline (AUC/AP) of the model after permutation and compare it with the original performance.

#### Significance level

Repeat the permutation N=1000 times and calculate the p-value of the original performance exceeding the permutation distribution (p<0.01 is considered significant).

All permutation scenarios lead to a significant performance decline, proving that the model relies on genuine biological features and topological structure.

The permutation test demonstrates that HyWinNet’s predictive capability stems from:

- **Feature-function association**: The true relationship between protein sequence/structure features and function (not random noise);
- **Topology-function coupling**: The hierarchical nature of PPI networks (e.g., modularity, scale-free properties) driving interactions;
- **Model interpretability**: Hyperbolic space embedding and graph wavelet transforms jointly extract biologically interpretable patterns, which are incompatible with the randomization hypothesis;

## 4. Discussion

Our study presents HyWinNet, a geometrically-principled deep learning framework that addresses fundamental limitations in current protein-protein interaction (PPI) prediction by integrating hyperbolic graph neural networks with multi-scale spectral analysis. The proposed model was motivated by the critical need to capture both hierarchical biological relationships and multi-resolution interaction patterns that are inherently present in protein networks but poorly modeled by conventional Euclidean-based approaches. Through systematic integration of Lorentzian hyperbolic embeddings, contrastive learning with feature augmentation, and graph wavelet transforms, HyWinNet demonstrates that geometric inductive biases combined with spectral signal processing can significantly enhance both the accuracy and interpretability of PPI predictions. The experimental results substantiate our core hypothesis that hyperbolic space representations coupled with multi-scale feature extraction would outperform existing methods in modeling complex biomolecular interactions, while providing biologically meaningful insights into interaction mechanisms at different structural resolutions.

HyWinNet achieves state-of-the-art performance (AUC: 0.922±0.005, AP: 0.938±0.005) in protein-protein interaction (PPI) prediction, outperforming six contemporary methods by 1.3-33.6%. These results align with but significantly extend prior work in three key aspects: (1) Like Struct2Graph (AUC: 0.892) and Fully_HNN (AUC: 0.909), we confirm that geometric deep learning architectures substantially improve PPI prediction over sequence-based methods (e.g., Topsy_Turvy’s AUC: 0.868). However, our integration of Lorentzian hyperbolic space with multi-scale wavelet transforms yields a 1.3-3.0% performance gain over these geometric approaches, suggesting that hierarchical embeddings alone are insufficient without explicit multi-resolution analysis. (2) While AGAT_PPIS and GACT_PPIS demonstrated the value of attention mechanisms (AUC: 0.866-0.878), our contrastive learning module with 20% feature dropout shows that view augmentation combined with hyperbolic distance metrics (Eq. 13-16) provides a more robust feature discrimination (AUC +4.4-5.6%). (3) Crucially, unlike purely Euclidean (GCN) or hyperbolic (HGCN) baselines, our ablation studies verify that the synergistic combination of Lorentz graph convolutions and multi-scale wavelet transforms (ΔAU C ≤0.0817 when removed) uniquely captures both hierarchical relationships and localized binding patterns. These differences likely stem from our biologically-aligned design: where prior works treated scale selection empirically (e.g., k=1-2 steps in Struct2Graph), our wavelet scales (k=1,2,3,4) explicitly model distinct biological resolutions from residue contacts to pathway interactions. This multi-scale geometric approach not only advances PPI prediction accuracy but also establishes a novel paradigm for interpretable analysis of hierarchical biological networks.

### 4.1. Data Set Analysis

Fig.12 depicts the distribution of the number of interactions per protein before and after applying data augmentation methods on the Luo and Zeng datasets. Both datasets (Luo and Zeng) demonstrate a broader distribution range after data augmentation. The augmentation effect is more pronounced in the Zeng dataset, manifested by a markedly extended distribution range, a substantial increase in the maximum number of interactions (with a more prominent upper extension), and a distinct upward shift in the median position. Data augmentation effectively elevates the density of the protein interaction network.

**Figure 12:**
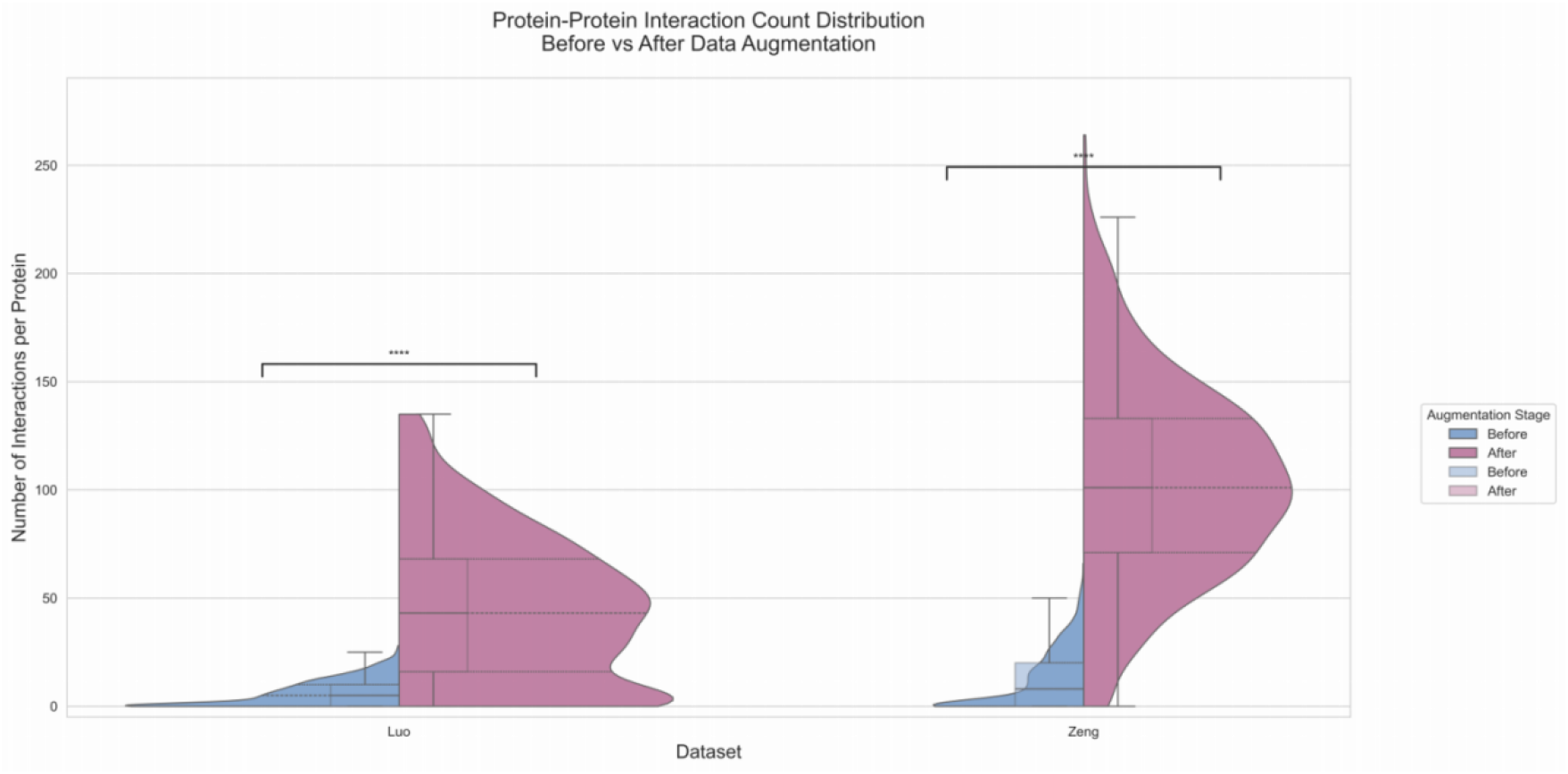
Comparison of protein-protein interaction frequencies before and after data augmentation across datasets

The data augmentation method employed herein is a graph data augmentation approach grounded in community detection. Its underlying principle involves leveraging the Louvain community detection algorithm to identify tightly connected modules (protein functional modules) within the network. Subsequently, new edges (interactions) are randomly introduced within the same module, thereby enhancing the internal connectivity of the module. This strategy preserves the topological structure of protein functional modules, and the augmented edges are more likely to mirror authentic biological relationships Fig.13 compares evaluation metrics and losses under diverse data augmentation methods. It is apparent that the graph data augmentation method based on community detection significantly outperforms other methods.

**Figure 13:**
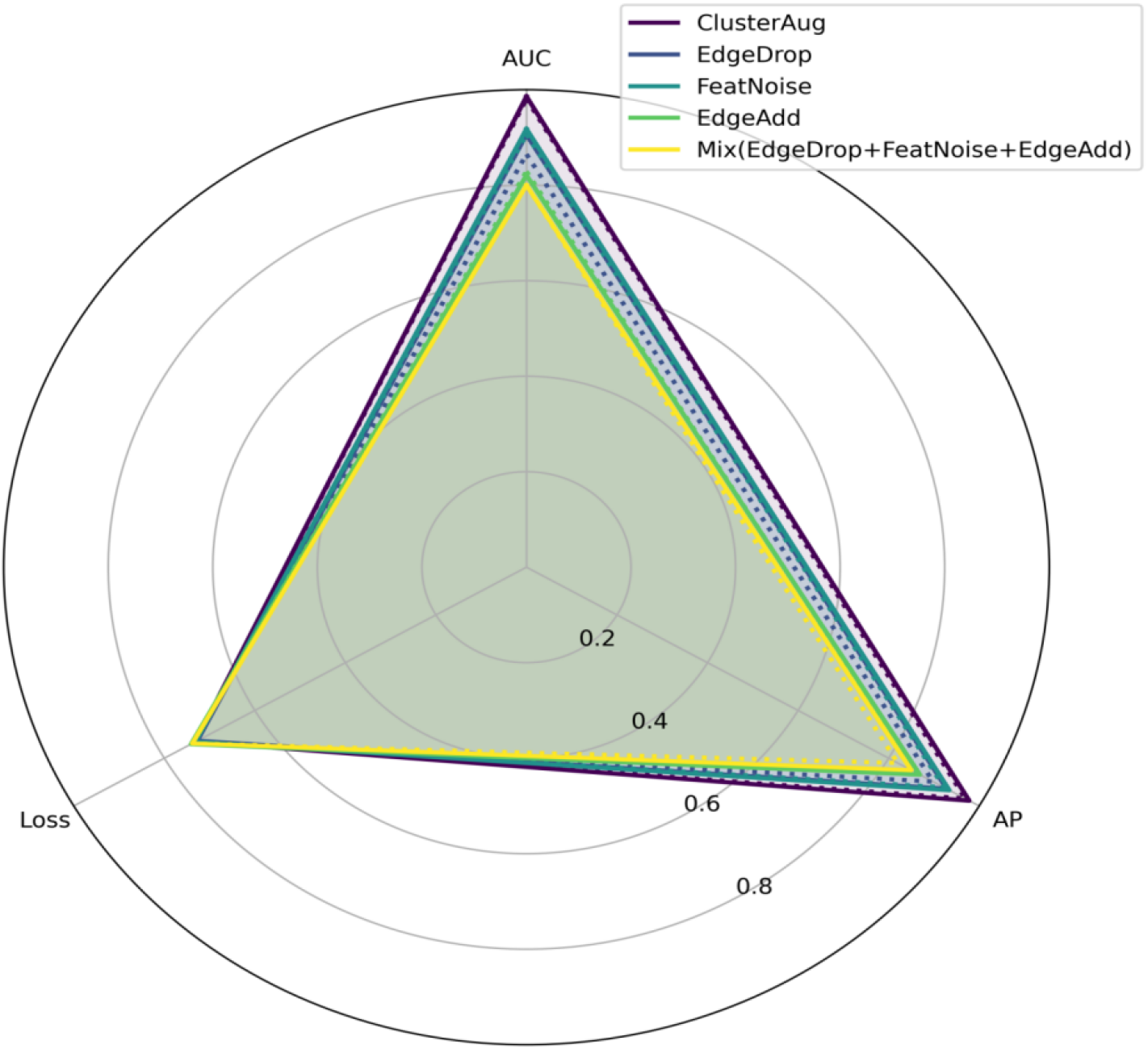
Comparison of evaluation metrics and loss across different data augmentation methods

### 4.2. Robustness Analysis

Hyperbolic spaces mitigate sensitivity to local perturbations through curvature-regulated global geometry[36], while multi-scale wavelet transforms synergistically integrate hierarchical information flow to prevent noise amplification inherent in single-scale representations. The multi-scale diffusion operator mitigates overfitting to local noise by integrating symmetric normalized with varying step sizes. Global-scale components (e.g., s=4) capture modular structures through long-range interactions, while local-scale counterparts (s=1) preserve proximal topological relationships – their synergistic combination inherently filters high-frequ-ency noise through spectral domain normalization. The above-mentioned measures collectively constitute a robust-ness-oriented design framework.

### 4.3. Generalization Analysis

Contrastive learning enforces semantic invariance (e.g., functional similarity) in protein interaction modeling, mitigating reliance on dataset-specific patterns, while Lorentzian geometric constraints regularize the learning process to prevent overfitting and improve generalization to unseen distributions[37].

### 4.4. Limitations

Although HyWinNet demonstrates superior performance in PPI prediction, three key limitations warrant consideration. First, the generalizability of the model is limited by the characteristics of the data set: Both the Luo et al.[26] and Zeng et al.[27] datasets predominantly cover human and model organism interactions, potentially limiting the applicability to understudied species or types of atypical interaction (e.g., transient or condition-specific PPIs). Second the current multi-scale wavelet transform operates on static graph structures, failing to fully capture the dynamic nature of protein interactions in vivo, where conformational changes and post-translational modifications critically influence binding events. Third, despite achieving state-of-the-art metrics (AUC: 0.922±0.005), the model’s interpretability remains partial: while hyperbolic embeddings provide geometric intuition for hierarchy-aware relationships, the biological meaning of specific wavelet coefficients (28) at different scales requires further experimental validation. These limitations highlight the need for future work incorporating temporal dynamics modeling and expanded taxonomic coverage in training data.

## Funding

This work was financially funded by National Natural Science Foundation of China (NSFC)(Project No. 62562041), the Science and Technology Project of Jiangxi Provincial Department of Education(Project No.GJJ2201043) and the university’s startup funding for new Ph.D.researchers (Project No.000/20298613).

## Declaration of Competing Interest

The authors declare that they have no known competing financial interests or personal relationships that could have appeared to influence the work reported in this paper.

## CRediT authorship contribution statement

**Qingzhi Yu**: Writing – original draft, Formal analysis, Conceptualization.. **Shuai Yan**: Writing – review & editing, Conceptualization.. **Wenfeng Dai**: lizatioWriting – review & editing, Conceptuan.. **Zhengrong Xi**: Visualization. **Yuxin Cheng**: Visualization. **Xiang Cheng**: Writing - review & editing, Supervision, Funding acquisition, Conceptualization.

## Data sources

The research utilizes two publicly accessible datasets for analysis. The primary dataset, initially reported in Nature Communications [26], consolidates three types of biomedical relationships: pharmaceutical compound-protein interactions, medication-pathology correlations, and inter-protein associations. This comprehensive resource can be accessed at: https://github.com/luoyunan/DTINet.

Additionally, a secondary dataset published in Chemical Science [27] was employed, featuring systematically organized data on compound-protein binding relationships extracted from diverse biomedical networks.This dataset is publicly available at: https://github.com/ChengF-Lab/deepDTnet.

## Notes

### Competing Interest Statement

The authors have declared no competing interest.

## References

[1] Jian Zhang and Lukasz Kurgan. Review and comparative assessment of sequence-based predictors of protein-binding residues. Briefings in bioinformatics, 19(5): 821–837, 2018.

[2] Preeti Thareja, Rajender Singh Chhillar, Sandeep Dalal, Sarita Simaiya, Umesh Kumar Lilhore, Roobaea Alroobaea, Majed Alsafyani, Abdullah M Baqasah, and Sultan Algarni. Intelligence model on sequence-based prediction of ppi using aisso deep concept with hyperparameter tuning process. Scientific Reports, 14(1): 21797, 2024.

[3] Ulrich Stelzl, Uwe Worm, Maciej Lalowski, Christian Haenig, Felix H Brembeck, Heike Goehler, Martin Stroedicke, Martina Zenkner, Anke Schoenherr, Susanne Koeppen, et al. A human protein-protein interaction network: a resource for annotating the proteome. Cell, 122(6):957–968, 2005.

[4] Bruce Alberts. The cell as a collection of protein machines: Preparing the next generation of molecular biologists. Cell, 92(3): 291–294, 1998.

[5] Wim Pomp, Joseph V.W. Meeussen, and Tineke L. Lenstra. Transcription factor exchange enables prolonged transcriptional bursts. Molecular Cell, 84(6):1036–1048.e9, 2024.

[6] Tony Pawson and Michael Kofler. Kinome signaling through regulated protein–protein interactions in normal and cancer cells. Current Opinion in Cell Biology, 21(2): 147–153, 2009. Cell regulation.

[7] Wei-Tse Hsu, Chi Nam Ignatius Pang, Josefa Sheetal, and Marc R. Wilkins. Protein–protein interactions and disease: Use of s. cerevisiae as a model system. Biochimica et Biophysica Acta (BBA) - Proteins and Proteomics, 1774(7): 838–847, 2007.

[8] Xiaopei Cui, Guopin Pan, Ye Chen, Xiaosun Guo, Tengfei Liu, Jing Zhang, Xiaofan Yang, Mei Cheng, Haiqing Gao, and Fan Jiang. The p53 pathway in vasculature revisited: A therapeutic target for pathological vascular remodeling? Pharmacological Research, 169:105683, 2021.

[9] Qiao Wang, Qinghe Li, Ranran Liu, Maiqing Zheng, Jie Wen, and Guiping Zhao. Host cell interactome of pa protein of h5n1 influenza a virus in chicken cells. Journal of Proteomics, 136: 48–54, 2016.

[10] Mike Boxem, Zoltan Maliga, Niels Klitgord, Na Li, Irma Lemmens, Miyeko Mana, Lorenzo de Lichtervelde, Joram D. Mul, Diederik van de Peut, Maxime Devos, Nicolas Simonis, Muhammed A. Yildirim, Murat Cokol, Huey-Ling Kao, Anne-Sophie de Smet, Haidong Wang, Anne-Lore Schlaitz, Tong Hao, Stuart Milstein, Changyu Fan, Mike Tipsword, Kevin Drew, Matilde Galli, Kahn Rhrissorrakrai, David Drechsel, Daphne Koller, Frederick P. Roth, Lilia M. Iakoucheva, A. Keith Dunker, Richard Bonneau, Kristin C. Gunsalus, David E. Hill, Fabio Piano, Jan Tavernier, Sander van den Heuvel, Anthony A. Hyman, and Marc Vidal. A protein domain-based interactome network for c. elegans early embryogenesis. Cell, 134(3): 534–545, 2008.

[11] Min Zeng, Fuhao Zhang, Fang Xiang Wu, Yaohang Li, and Min Li. Protein-protein interaction site prediction through combining local and global features with deep neural networks. Bioinformatics, 36(4), 2019.

[12] Tristan T Aumentado-Armstrong, Bogdan Istrate, and Robert A Murgita. Algorithmic approaches to protein-protein interaction site prediction. Algorithms for Molecular Biology, 10(1):7, 2015

[13] Jack F. Greenblatt, Bruce M. Alberts, and Nevan J. Krogan. Discovery and significance of protein-protein interactions in health and disease. Cell, 187(23): 6501–6517, 2024.

[14] Wenxing Hu and Masahito Ohue. Spatialppiv2: Enhancing protein–protein interaction prediction through graph neural networks with protein language models. Computational and Structural Biotechnology Journal, 27: 508–518, 2025.

[15] Fan Zhang, Sheng Chang, Binjie Wang, and Xinhong Zhang. Dssgnn-ppi: A protein–protein interactions prediction model based on double structure and sequence graph neural networks. Computers in Biology and Medicine, 177:108669, 2024.

[16] Kamal Taha. Protein-protein interaction detection using deep learning: A survey, comparative analysis, and experimental evaluation. Computers in Biology and Medicine, 185:109449, 2025.

[17] Li Yiwei, Golding G Brian, and Ilie Lucian. Delphi: accurate deep ensemble model for protein interaction sites prediction. Bioinformatics (Oxford, England), 37(7): 896–904, 2020.

[18] Wenxing Hu and Masahito Ohue. Spatialppi: Three-dimensional space protein-protein interaction prediction with alphafold multimer. Computational and Structural Biotechnology Journal, 23: 1214–1225, 2024.

[19] Wenjian Ma, Xiangpeng Bi, Huasen Jiang, Zhiqiang Wei, and Shugang Zhang. Annotating protein functions via fusing multiple biological modalities. Communications Biology, 7(1): 1705–1705, 2024.

[20] Xin Zeng, Fan-Fang Meng, Xin Li, Kai-Yang Zhong, Bei Jiang, and Yi Li. Ghgpr-ppis: A graph convolutional network for identifying protein-protein interaction site using heat kernel with generalized pagerank techniques and edge self-attention feature processing block. Computers in Biology and Medicine, 168:107683, 2024.

[21] Xiaohan Sun, Zhixiang Wu, Jingjie Su, and Chunhua Li. Graphpbsp: Protein binding site prediction based on graph attention network and pre-trained model prostt5. International Journal of Biological Macromolecules, 282:136933, 2024.

[22] Lu Meng, Lishuai Wei, and Rina Wu. Mvgnn-ppis: A novel multi-view graph neural network for protein-protein interaction sites prediction based on alphafold3-predicted structures and transfer learning. International Journal of Biological Macromolecules, 300:140096, 2025.

[23] Ziqi Gao, Chenran Jiang, Jiawen Zhang, Xiaosen Jiang, Lanqing Li, Peilin Zhao, Huanming Yang, Yong Huang, and Jia Li. Hierarchical graph learning for protein–protein interaction. Nature Communications, 14(1): 1093, 2023.

[24] Seungsik Min, Kyungsik Kim, Ki-Ho Chang, Deok-Ho Ha, and Jun-Ho Lee. Topological properties of four networks in protein structures. Physica A: Statistical Mechanics and its Applications, 486: 956–967, 2017.

[25] Subhankar Sarkar and Souvik Chakraborty. Spatio-spectral graph neural operator for solving computational mechanics problems on irregular domain and unstructured grid. Computer Methods in Applied Mechanics and Engineering, 435:117659, 2025.

[26] Yunan Luo, Xinbin Zhao, Jingtian Zhou, Jinglin Yang, Yanqing Zhang, Wenhua Kuang, Jian Peng, Ligong Chen, and Jianyang Zeng. A network integration approach for drug-target interaction prediction and computational drug repositioning from heterogeneous information. Nature Communications, 8(1): 573, 2017.

[27] Xiangxiang Zeng, Siyi Zhu, Weiqiang Lu, Zehui Liu, Jin Huang, Yadi Zhou, Jiansong Fang, Yin Huang, Huimin Guo, Lang Li, Bruce D. Trapp, Ruth Nussinov, Charis Eng, Joseph Loscalzo, and Feixiong Cheng. Target identification among known drugs by deep learning from heterogeneous networks electronic supplementary information (esi) available. see doi: 10.1039/c9sc04336e. Chemical Science, 11(7):1775–1797, 2020.

[28] Weize Chen, Xu Han, Yankai Lin, Hexu Zhao, Zhiyuan Liu, Peng Li, Maosong Sun, and Jie Zhou. Fully hyperbolic neural networks. 2021.

[29] Siddharth Viswanath, Hiren Madhu, Dhananjay Bhaskar, Jake Kovalic, Dave Johnson, Rex Ying, Christopher Tape, Ian Adelstein, Michael Perlmutter, and Smita Krishnaswamy. Hiponet: A topology-preserving multi-view neural network for high dimensional point cloud and single-cell data. 2025.

[30] Thomas Villmann, Marika Kaden, Mandy Lange, Paul Sturmer, and Wieland Hermann. Precision-recall-optimization in learning vector quantization classifiers for improved medical classification systems. In Computational Intelligence & Data Mining, 2014.

[31] Mei-Yuan Cao, Suhaila Zainudin, and Kauthar Mohd Daud. Feature fusion with attributed deepwalk for protein–protein interaction prediction. Scientific Reports, 15(1): 12255, 2025.

[32] Zhou Yuting, Jiang Yongquan, and Yang Yan. Agat-ppis: a novel protein-protein interaction site predictor based on augmented graph attention network with initial residual and identity mapping. Briefings in bioinformatics, 24(3), 2023.

[33] Lu Meng and Huashuai Zhang. Gact-ppis: Prediction of protein-protein interaction sites based on graph structure and transformer network. International Journal of Biological Macromolecules, 283(P1):137272–137272, 2024.

[34] Baranwal Mayank, Magner Abram, Saldinger Jacob, Turali Emre Emine S., Elvati Paolo, Kozarekar Shivani, VanEpps J. Scott, Kotov Nicholas A., Violi Angela, and Hero Alfred O. Struct2graph: a graph attention network for structure based predictions of protein–protein interactions. BMC Bioinformatics, 23(1): 370–370, 2022.

[35] Singh Rohit, Devkota Kapil, Sledzieski Samuel, Berger Bonnie, and Cowen Lenore. Topsy-turvy: integrating a global view into sequence-based ppi prediction. Bioinformatics (Oxford, England), 38(Supplement_1):i264–i272, 2022.

[36] Huijuan Hu, Chaobo He, Xinran Chen, and Quanlong Guan. Hckgl: Hyperbolic collaborative knowledge graph learning for recommendation. Neurocomputing, 634:129808, 2025.

[37] Shengrui Xu, Tianchi Lu, Zikun Wang, Jixiu Zhai, and Jingwan Wang. Scmppi: Supervised contrastive multimodal framework for predicting protein-protein interactions. 2025.

